# De novo genome assembly of Ansell’s mole-rat (*Fukomys anselli*)

**DOI:** 10.1101/2025.06.03.657636

**Authors:** Milica Bekavac, Raphael Coimbra, Veronica F. Busa, Mikaela Behm, Rebecca E. Wagner, Angela Goncalves, Sabine Begall, Michaela Frye, Duncan T. Odom

**Author notes:** Co-first author (Milica Bekavac, Raphael Coimbra, Veronica F. Busa). Co-corresponding &.

## Abstract

Ansell’s mole-rat (*Fukomys anselli*) is an African rodent known for its subterranean lifestyle and unique phenotypic traits, including extreme longevity, magnetoreception, and a cooperative breeding social structure. Efforts to dissect the genetic architecture of these traits and to decipher their phylogenetic relationships within the broader African mole-rat family would greatly benefit from a reference-grade genome. Here, we report a first genome assembly of a male Ansell’s mole-rat. By combining Oxford Nanopore Technologies (ONT) long-reads and Illumina short-reads with Hi-C data, we generated a chromosome level assembly with a total length of 2.27 Gb, 412 scaffolds and a scaffold N50 of 72.4 Mb. We identified 99.54% of expected genes and annotated 29,094 transcripts using RNA sequencing data. This high-quality *de novo* genome of *Fukomys anselli* lays the foundation for dissecting the genetic and evolutionary basis of its extraordinary traits and resolving African mole-rat phylogeny.

**ARTICLE SUMMARY:** Bekavac, Coimbra, and Busa *et al*. assemble a *de novo* genome of a male *Fukomys anselli* individual. They use short-read and long-read whole genome sequencing and Hi-C sequencing to achieve chromosome-level contiguity, and annotate the genome using RNA-seq from nine diverse tissues. This high-quality genome enables exploration of the genetic basis of Ansell’s mole-rat’s remarkable traits, including longevity, hypoxia tolerance, and magnetoreception. It also allows for comprehensive analysis of the currently-contested phylogenetic relationships within the mole-rat family.

## INTRODUCTION

Ansell’s mole-rat (*Fukomys anselli*) is a subterranean rodent endemic to Zambia (Burda et al. 1999). Like other species within the African mole-rats family (Bathyergidae), Ansell’s mole-rat lives in complex, narrow, self-excavated tunnels and has developed multiple morphological and physiological adaptations to this underground lifestyle (reviewed in Begall et al. 2021). The animals tolerate low levels of O_2_ / high levels of CO_2_ and respond to hypoxia / hypercapnia with downregulation of free thyroid hormone T3 and erythropoiesis (Henning et al. 2024). Ansell’s mole-rat compensates for reduced visual (Němec et al. 2004; Wegner et al. 2006a) and hearing capabilities (Gerhardt et al. 2017) with a well-developed olfactory system (Caspar et al. 2022) and magnetoreception (Burda et al. 1990; Marhold et al. 1997; Wegner et al. 2006b; Caspar et al. 2020) to navigate through tunnels.

Ansell’s mole-rats are exceptionally long-lived among rodents, with adult lifespans up to 22 years (Dammann et al. 2022). They live in cooperative breeding groups with one breeding pair and non-breeding offspring (Patzenhauerová et al. 2013). Interestingly, lifespan within the *Fukomys* genus is notably shorter for non-breeders compared to breeders, providing an opportunity to study gene regulation and expression that affects aging between closely-related individuals (Dammann et al. 2011; Sahm et al. 2021; Dammann et al. 2023). However, current research efforts in the molecular mechanisms underlying longevity and social structures have been limited by a lack of high-resolution genomes for comparative studies within the *Fukomys* genus (Sahm et al. 2021).

Extreme longevity is not unique to *F. anselli* within the Bathyergidae family. Notable examples of exceptionally long lifespan include the naked mole-rat (*Heterocephalus glaber*, maximum age 40 years) and Damaraland mole-rat (*Fukomys damarensis*, maximum age > 20 years) (Fang et al. 2014; Dammann et al. 2019; Buffenstein and Amoroso 2024; Ruby et al. 2024). These species have become important resources for research in cancer (Liang et al. 2010; Tian et al. 2013), hypoxia tolerance (Park et al. 2017), pain insensitivity (Park et al. 2008), and reproduction (Schmidt et al. 2014; Brieño-Enríquez et al. 2023). The genomes of *H. glaber* and *F. damarensis* are the only available assemblies for the Bathyergidae family (Fang et al. 2014; Keane et al. 2014; Sokolowski et al. 2024).

Comparative genomics efforts resting on *de novo* transcriptomes have been used to analyse Bathyergidae divergence (Davies et al. 2015; Sahm, Bens, Szafranski, et al. 2018; Sahm et al. 2021). However, these approaches are limited by: input data quality (Smith-Unna et al. 2016), a reliance on only the longest isoform for expression quantification (Davies et al. 2015; Sahm, Bens, Henning, et al. 2018; Sahm et al. 2021), unreliable expression estimates due to fragmented contiguous sequences (Hsieh et al. 2019) or short transcript lengths (Wu et al. 2018), incomplete transcript assembly (Ungaro et al. 2017), and species-specific performance of assemblers (Hölzer and Marz 2019). Harnessing high-quality whole-genome assemblies from multiple Bathyergidae species empowers comprehensive interspecies analyses and eliminates the barriers imposed by transcriptome-only comparisons.

Ansell’s mole-rat was first formally described in Burda et al. 1999 as *Cryptomys anselli*, with a distinguishing karyotype of 2n=68. Further phylogenetic and karyotypic evidence led to the taxonomic separation of *Fukomys* from the *Cryptomys* genus (Kock et al. 2006). Nevertheless, the Bathyergidae family phylogeny remains debated (Visser et al. 2019) (see below). Prior efforts have leveraged 12S rRNA, TTR intron I, and mitochondrial cytb sequencing to genetically resolve phylogenetic relationships (Faulkes et al. 2004; Ingram et al. 2004; Van Daele et al. 2007). To comprehensively analyse phylogenetic relationships within the mole-rat family will require highest quality whole genome assemblies, thus motivating our efforts to generate a full *F. anselli* genome.

Here, we report the first chromosome-scale genome of a male *Fukomys anselli*, assembled using ONT, Illumina, and Hi-C sequencing. This high-quality genome provides the foundation for uncovering the genetic basis of Ansell’s mole-rat’s remarkable traits, including longevity, hypoxia tolerance, and magnetoreception.

## METHODS AND MATERIALS

### Sample collection

Animals were housed at the University of Duisburg-Essen (approved by permit no. 32-2-1180-71/328 Veterinary Office of the City of Essen) in a humidity and temperature-controlled room with a 12L:12D cycle. Ambient temperature was kept constant at 24 ± 1 °C with relative humidity of 40-50 %. The animals were fed with raw carrots and potatoes three times per week as staple food, supplemented with cereals and apples once per week and lettuce on an irregular basis. No free water was needed. They were kept in family groups in glass terraria ranging in size (W x L x H) from 45 cm x 70 cm x 40 cm to 60 cm x 140 cm x 40 cm, depending on family size. The terraria were filled with sawdust, and plastic or wooden tubes enriched the terraria. Flower pots served as nests, hay and paper strips were offered as nesting material.

The animals were put under deep anesthesia with an intramuscular injection of 12 mg/kg ketamine (Ceva GmbH) and 5 mg/kg xylazine (Ceva GmbH) and then decapitated. We collected organs for long-read sequencing, short-read sequencing, Hi-C sequencing and RNA sequencing from a 21-month-old male. Liver, spleen, testes, calf muscle, kidney, heart, lung were dissected and flash frozen. In addition, flash frozen brain and skin from two other males, 12- and 19-month-old individuals, respectively, were also used for RNA sequencing experiments.

### Nanopore long-read sequencing

Frozen liver was pulverized using CP02 CryoPrep Automated Dry Pulverizer (Covaris) and genomic DNA was isolated using the Puregene® Tissue kit (Qiagen) according to manufacturer’s instructions, with the following modifications in case of using the ONT Ultra-Long DNA Sequencing Kit V14 for subsequent library preparation: the final elution buffer was changed to EEB buffer from Oxford Nanopore Technologies (ONT) and the first incubation for dissolving DNA decreased to 56°C for 20 minutes. One DNA extraction was performed using the Nanobind® PanDNA kit (PacBio), according to the manufacturer’s instructions, with a change in the initial tissue processing (CryoPrep to disrupt the tissue) and the final elution buffer to EEB buffer (ONT). The DNA was quantified using the Qubit™ dsDNA BR Assay kit (Invitrogen) and the DNA integrity was verified with Genomic DNA ScreenTape Assay on a TapeStation (Agilent Technologies). Libraries were prepared using the ONT Ultra-Long DNA Sequencing Kit V14 (SQK-ULK114) (starting from the Tagmentation reaction) or ONT Ligation Sequencing kit V14 (SQK-LSK114). Sequencing was performed on one MinION flow cell on a GridION instrument and 7 PromethION flow cells (R10.4.1), which generated a total of 157.41 GBases.

### Illumina short-read sequencing

DNA was isolated from a frozen spleen fragment using AllPrep® DNA/RNA/Protein Mini kit (Qiagen) according to the manufacturer’s instructions. The DNA was quantified using the Qubit™ dsDNA BR Assay kit (Invitrogen) and its integrity was verified with Genomic DNA ScreenTape Assay on a TapeStation (Agilent Technologies). The library was prepared using the Illumina DNA PCR-Free Library Prep kit, according to the manufacturer’s instructions. It was quantified with Qubit™ ssDNA Assay Kit (Invitrogen) and sequenced on Illumina NovaSeq X Plus instrument in a 2×150 bp configuration, which generated 400.81 GBases of data.

### Hi-C sequencing

Arima High Coverage HiC kit (Arima Genomics) was used to generate proximally-ligated DNA according to the manufacturer’s protocol (A160162 v01), with the following modifications: the liver was pulverized with CP02 CryoPrep Dry Pulverizer (Covaris) instead of mortar and pestle; step 2 of the proximal ligation was extended to 20 minutes. A library compatible with Illumina sequencing was prepared with the Arima Library Prep Module (Arima Genomics), according to the protocol for Arima High Coverage HiC kit (Document number: A160186 v02). The concentration and size distribution of the library were measured with Qubit™ 1X dsDNA High Sensitivity Assay kit (Invitrogen) and D5000 ScreenTape Assay on TapeStation (Agilent Technologies), respectively. The library was sequenced on two lanes of Illumina NovaSeq 6000 instrument in 2×250 bp mode. In total, 355.93 GBases of data was generated.

### RNA sequencing

Liver, spleen, testes, calf muscle, kidney, heart, lung and brain were pulverized using CP02 CryoPrep Dry Pulverizer (Covaris). The tissue powder was placed in tubes with metal beads and TRIzol™ (Invitrogen) and homogenized with TissueLyser (Qiagen). The debris was removed by centrifugation and total RNA was extracted with Direct-zol™ RNA MiniPrep kit (Zymo Research) according to manufacturer’s instructions. The RNA quality was assessed with High Sensitivity RNA ScreenTape Assay on a TapeStation (Agilent Technologies) and quantity was measured with Qubit™ RNA BR Assay kit (Invitrogen). Libraries were prepared using the Illumina Stranded Total RNA Prep, Ligation with Ribo-Zero Plus kit according to the manufacturer’s instructions. The quality of the libraries was assessed on a D5000 ScreenTape Assay on a TapeStation (Agilent Technologies) and their concentration was measured with Qubit™ 1X dsDNA High Sensitivity Assay kit (Invitrogen). Pooled libraries were sequenced on the Illumina NovaSeq X Plus instrument in 2×100 bp mode.

For skin RNA-seq, total RNA was extracted from abdomen skin tissue of a single *Fukomys anselli* individual using a combination of TRIzol™ (Invitrogen) and the mirVana™ miRNA Isolation Kit (Thermo Fisher Scientific). RNA quantity and quality were assessed using the Qubit™ RNA BR Assay Kit (Invitrogen) and the Bioanalyzer RNA 6000 Nano Assay (Agilent Technologies). To eliminate genomic DNA contamination, the RNA was treated with TURBO DNase (Invitrogen), followed by purification using the RNA Clean & Concentrator™ kit (Zymo Research). RNA-seq libraries were prepared using the TruSeq Stranded Total RNA Library Prep Kit (Illumina) with IDT for Illumina TruSeq DNA/RNA Unique Dual Indexes (UDI). Sequencing was carried out on one Illumina NextSeq 550 lane using a 150 bp paired-end configuration.

### Assembly

Prior to assembly, we estimated the genome size and repeat coverage based on *k*-mer frequencies in the short-read sequencing via Jellyfish v2.3.1 and GenomeScope v2.0.1 (Marçais and Kingsford 2011; Ranallo-Benavidez et al. 2020). We base-called the long-read sequencing using the sup model of Dorado v.0.9.0 (Oxford Nanopore Technologies PLC. Public License, v. 1.0), then assembled and performed two rounds of polishing on the long-read sequencing via Flye v2.9.5-b1801 (Kolmogorov et al. 2019) using a GenomeScope-based genome size estimate of 2.3 Gb and the argument –nano-hq. Although we attempted further polishing using medaka v2.0.1, this was detrimental to genome assembly based on multiple quality assessment metrics such as error rate, *k*-mer completeness, contig N50, and gene completeness, so this additional polishing was omitted in the final version. We leveraged the short-read sequencing to correct base errors (SNVs/indels) in the polished assembly via NextPolish v1.4.1 (Hu et al. 2020).

We then used purge_dups v1.2.6 (Guan et al. 2020) to remove haplotigs and overlaps. Only haplotigs and overlaps at the end of contigs were selected for removal to avoid omitting false positive duplications in the middle of contigs. We used the BLAST+ v2.16.0 executable blastn (Camacho et al. 2009) to identify possible contamination, minimap2 v2.28 (Li 2021) and samtools v1.21 (Danecek et al. 2021) to generate duplicate-marked short-read alignments to the purged assembly, and BlobToolkit v4.4.0 (Challis et al. 2020) to calculate and visualize contig coverage and GC content. Contigs were removed as contamination if they returned no hits (6715 contigs) or were assigned to orders other than Rodentia and did not match any known mole-rat sequences (8 contigs). Additionally, all contigs shorter than 1 kbp (182 contigs) were removed prior to scaffolding.

Hi-C sequencing was cleaned via fastp v0.24.0 (Chen et al. 2018) to trim adapters and polyG sequencing artifacts and to discard low-quality reads and reads shorter than 36 bp, then processed and aligned to the cleaned contigs via a modified Arima Genomics mapping pipeline (doc A160156 v03 January 2024). Hi-C scaffolding was performed using YaHS v1.2.2 (Zhou et al. 2023), and the assembly was manually curated for misjoins, missed joins, translocations, and inversions using Juicebox with Assembly Tools v2.17.0 (Durand et al. 2016; Dudchenko et al. 2018). The final, curated assembly was visualized for synteny to *Fukomys damarensis* (DMR_v1.0_HiC) via JupiterPlot v1.1 (Chu 2018).

### Quality assessment

We quantified the quality of our multiple long-read sequencing runs using NanoComp v1.24.2 and NanoPlot v1.43.0 (De Coster and Rademakers 2023) (**Figure S1a, Table S1**).

We employed QUAST v5.3.0 (Gurevich et al. 2013) to measure assembly contiguity, Mercury with Meryl v1.4.1 (Rhie et al. 2020) to measure consensus quality (QV), error rate, and *k*-mer completeness, and compleasm v0.2.6 (Huang and Li 2023) to measure assembly completeness based on the “glires_odb10” gene set at all steps of the assembly pipeline to track assembly progress. We also used these tools to compare our *de novo* Ansell’s mole-rat assembly to other available rodent genomes.

Short-read RNA-seq of all nine tissues was processed via fastp v0.24.0 (Chen et al. 2018) to trim adapters and polyG sequencing artifacts and to discard low-quality reads and reads shorter than 36 bp (**Figure S1b**). FastQ Screen v0.16.0 (Wingett and Andrews 2018) was used to validate that the library predominantly aligns with the available *Fukomys damarensis* genome rather than potential contaminant genomes (**Figure S1c**). MultiQC v1.27.1 (Ewels et al. 2016) was used to visualise the RNA-seq quality.

OMArk v0.3.1 (Nevers et al. 2025) provided a proteome completeness and consistency assessment.

### Annotation

Repeats and transposable elements were identified and annotated using Earl Grey v5.1.1 (Baril et al. 2024) based on a *de novo* repeat library and known *Fukomys damarensis* repeats from the Dfam database v3.7 (Hubley et al. 2016). Identified repeats were soft masked prior to gene annotation using Earl Grey.

Short-read RNA-seq derived from nine different tissues was aligned to the assembled scaffolds using STAR v2.7.11b (Dobin et al. 2013) in two-pass mode for splice-aware mapping. We then applied BRAKER v3.0.8 (Hoff et al. 2019; Gabriel et al. 2024) for structural gene annotation using the aligned RNA-seq along with protein sequences derived from OrthoDB v12 vertebrates (Tegenfeldt et al. 2025) and from 41 NCBI RefSeq annotated rodent species (**Table S2**). Gene identification was optimized by assigning the argument busco_lineage to glires_odb10.

### De novo assembly and annotation of the mitochondrial genome

The mitogenome was assembled using the short-read sequencing via GetOrganelle v1.7.7.1 (Jin et al. 2020) using the *Fukomys damarensis* (NC_027742.1) and *Heterocephalus glaber* (NC_015112.1) mitogenomes as seeds. Next, we applied MitoAnnotator v4.0.9 (Zhu et al. 2023) to annotate the newly assembled mitogenome. The annotated mitochondrial features were visually inspected and start/end positions were corrected when necessary based on comparisons with *F. damarensis* and *H. glaber* mitochondrial gene sequences and annotations. Modified OrganellarGenomeDRAW v1.3.1 (Greiner et al. 2019) outputs were used for visualization.

To confirm the mitogenome assembly using long-read ONT sequencing, potential mitochondrial reads were first extracted by mapping to *F. damarensis* mitogenome via minimap2 v2.28 (Li 2021). Reads shorter than 1 kb or longer than 20 kb were excluded. The remaining reads were compared to the *F. damarensis* reference mitogenome using BLAST+ v2.16.0 (Camacho et al. 2009) and were retained only if they covered at least 70% of the reference, normalized by the subject (read) length. We assembled and polished the mitogenome from filtered reads via Flye v2.9.5-b1801 (Kolmogorov et al. 2019); this resulted in a single, circular contig. Error correction was performed using short reads trimmed with fastp v0.24.0 (Chen et al. 2018) via Pilon v1.24 (Walker et al. 2014).

To verify the identity of our mitogenome, the final assembly sequence was aligned to four publicly available 12S rRNA sequences from different *F. anselli* individuals (AY427022-AY427025) (Ingram et al. 2004) using the L-INS-i algorithm in MAFFT v7.526 (Katoh and Standley 2013).

## RESULTS AND DISCUSSION

We *de novo* assembled and annotated the genome of a male Ansell’s mole-rat (*Fukomys anselli*) using multiple modalities. Briefly, we performed Oxford Nanopore long-read sequencing (ONT) using DNA isolated from liver to obtain a whole-genome coverage of 46x and Illumina short-read sequencing (WGS) using DNA isolated from spleen to obtain a whole-genome coverage of >175x. We used Flye to assemble the ONT, and NextPolish to correct base errors in the sequence using WGS (Kolmogorov et al. 2019; Hu et al. 2020). We then cleaned contigs using purge_dups to remove haplotigs and BLASTn to identify and remove potential contamination (**Figure S2a**) (Camacho et al. 2009; Guan et al. 2020). This cumulatively produced a 2.41 Gb genome across 7581 contigs with an N50 of 19.8 Mb (**Figure S2b, Table S3**).

Sequence alignment of Hi-C data yielded 13.3 million unique inter-contig reads (**Table S4)**, which were leveraged by YaHS to create a scaffolded genome that was manually curated using Juicebox and Assembly Tools (**Figure 1a**, **Figure S2c**) (Durand et al. 2016; Zhou et al. 2023). The final *de novo* Ansell’s mole-rat assembly is 2.27 Gb across 412 scaffolds with an N50 of 72.4 Mb and L50 of 12 (**Figure 1b**, **Table 1**). This genome length is similar to expectations based on the 2.23 Gb estimated from *k*-mers by GenomeScope (Ranallo-Benavidez et al. 2020) and a previously-assembled 2.3 Gb *F. damarensis* genome (DMR_v1.0_HiC) (Fang et al. 2014).

**Figure 1:**
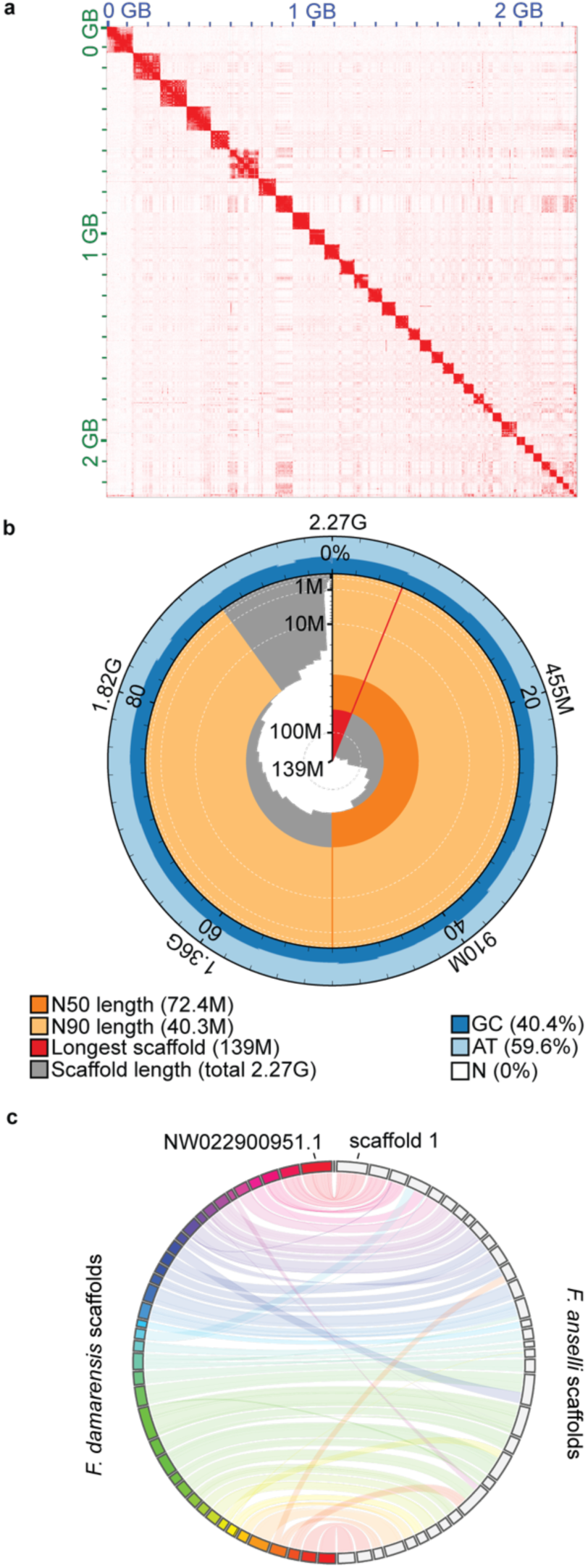
Chromosome-level *Fukomys anselli* genome assembly. a) Hi-C read contact map following final scaffold curation. The saturation of red corresponds to the number of contacts. b) Snail plot depicting the length and GC content of scaffolds. c) Circos plot depicting synteny between *Fukomys damarensis* (left, colorful) and *Fukomys anselli* (right, grey) scaffolds for all *F. damarensis* scaffolds longer than 10Mb and for *F. anselli* scaffolds constituting 99% of the genome. A fully annotated circos plot with scaffold numbers can be found in Figure S3.

**Table 1:** Contiguity of available mole-rat genomes.

|  | *F. anelli*<br>(This assembly) | *F. damarensis*<br>(DMR_v1.0_HiC) | *H. glaber*<br>(HetGla_female_1.0) | *H. glaber*<br>(mHetGlaV3) |
| --- | --- | --- | --- | --- |
| # contigs | 412 | 73,969 | 4,229 | 75 |
| # contigs<br>≥10 Mbp | 34 | 39 | 88 | 30 |
| Total length | 2,274,106,617 | 2,334,375,022 | 2,618,204,639 | 2,474,750,198 |
| N50 | 72,399,386 | 62,586,000 | 20,532,749 | 97,315,161 |
| N90 | 40,322,336 | 37,171,693 | 5,125,547 | 53,093,233 |
| auN | 80,188,288 | 66,455,256.8 | 22,553,264.2 | 94,268,059.9 |
| L50 | 12 | 15 | 42 | 11 |
| L90 | 28 | 34 | 142 | 25 |
| GC (%) | 40.43 | 40.34 | 40.21 | 39.96 |
| N's / 100 kbp | 1.45 | 2,072.24 | 11,589.37 | 71.12 |

Our highly contiguous *F. anselli* genome is the first chromosome-level *Fukomys* genome and demonstrates ortholog completeness superior to other currently-available resources. Gene analysis using compleasm (Huang and Li 2023) identified 99.54% ortholog completeness within the glires lineage, which represents all lagomorphs and rodents (**Table 2**). Prior research in the *Fukomys* genus has relied on incomplete and extremely fragmented genomes; for instance, *F. mechowii* and *F. micklemi* have only been analysed using short-read RNA-seq-derived *de novo* transcriptomes, both of which contained approximately 20,000 contigs with assembly completeness of 93.7% and 93.4%, respectively, estimated by BUSCO (Sahm, Bens, Szafranski, et al. 2018). Although there is an available Hi-C-scaffolded *F. damarensis* genome, the best published resource is composed of 73,969 scaffolds with a completeness estimate of 96.27% (**Table 1**, **Table 2**).

**Table 2:** Completeness estimates of available mole-rat genomes.

|  | *F. anelli*<br>(This assembly) | *F. damarensis*<br>(DMR_v1.0_HiC) | *H. glaber*<br>(HetGla_female_1.0) | *H. glaber*<br>(mHetGlaV3) |
| --- | --- | --- | --- | --- |
| Single-copy<br>complete | 99.09%, 13673 | 95.88%, 13230 | 97.59%, 13466 | 96.42%, 13304 |
| Duplicated<br>complete | 0.45%, 62 | 0.39%, 54 | 1.40%, 193 | 1.33%, 183 |
| Fragmented | 0.10%, 14 | 1.96%, 270 | 0.34%, 47 | 0.33%, 45 |
| Missing | 0.36%, 49 | 1.77%, 244 | 0.67%, 92 | 1.93%, 266 |

As expected, *F. anselli* demonstrates a high degree of synteny with *F. damarensis* (Van Daele et al. 2007): the 34 largest *F. anselli* scaffolds (each larger than 10 Mb) represent 99% of the genome and correspond to all 39 *F. damarensis* scaffolds larger than 10 Mb (**Figure 1c, Figure S3**). Although some Damaraland scaffolds are split across Ansell’s scaffolds and vice-versa (e.g. the intersecting bands in Figure 1c), there are few non-syntenic regions. Synteny with the *F. damarensis* scaffold NW_022900951.1, which is annotated for known X-linked genes such as Ar, Dmd, Fmr1, and Mecp2, suggests that the largest scaffold in the *F. anselli* genome, scaffold 1, is the X chromosome (**Figure 1c, Figure S3**). This is corroborated by the scaffold’s depleted sequencing coverage, consistent with X chromosome hemizygosity in our male individual (**Figure S2d**). There have been reports that the chromosomal complement of *F. anselli* – in particular the sex chromosomes – may be more complex than could be captured by our single-individual reference XY genome (Burda et al. 1999). Although all karyotyped *F. anselli* individuals have been found to have a diploid chromosome number of 2n=68, there appears to be variation in the size and centromeric positioning among both the sex and autosomal chromosomes. Indeed, the two X chromosomes in female karyotypes are often heteromorphic. Thus, to fully capture the standing structural variation among Ansell’s mole-rat may require sequencing of additional individuals of both sexes.

### Mitochondrial genome

We assembled the mitochondrial genome using GetOrganelle based on *H. glaber* and *F. damarensis* mitochondrial assemblies and annotated it using MitoAnnotator (Jin et al. 2020; Zhu et al. 2023). Consistent with other annotated mole-rat genomes, we identified the control region, 13 genes, two rRNAs, and 22 tRNAs in the 17,006 bp *F. anselli* mitogenome (**Figure 2**). The Ansell mole-rat’s mitogenome is larger than that of other closely-related species: 16,372 bp in *F. damarensis* and 16,386 bp in *H. glaber.* This difference is mostly attributable to a longer control region. To verify that this discrepancy is not attributable to artifacts of the assembly method, we independently assembled the mitogenome from ONT reads (see Methods), which generated a sequence differing by only one nucleotide (data not shown). Moreover, the alignment of the assembled mitogenome’s 12S rRNA region with four publicly available *F. anselli* 12S rRNA sequences revealed high sequence identity (Ingram et al. 2004). The alignments contained no gaps and few nucleotide differences, which are consistent with expected intraspecific variation (data not shown).

**Figure 2:**
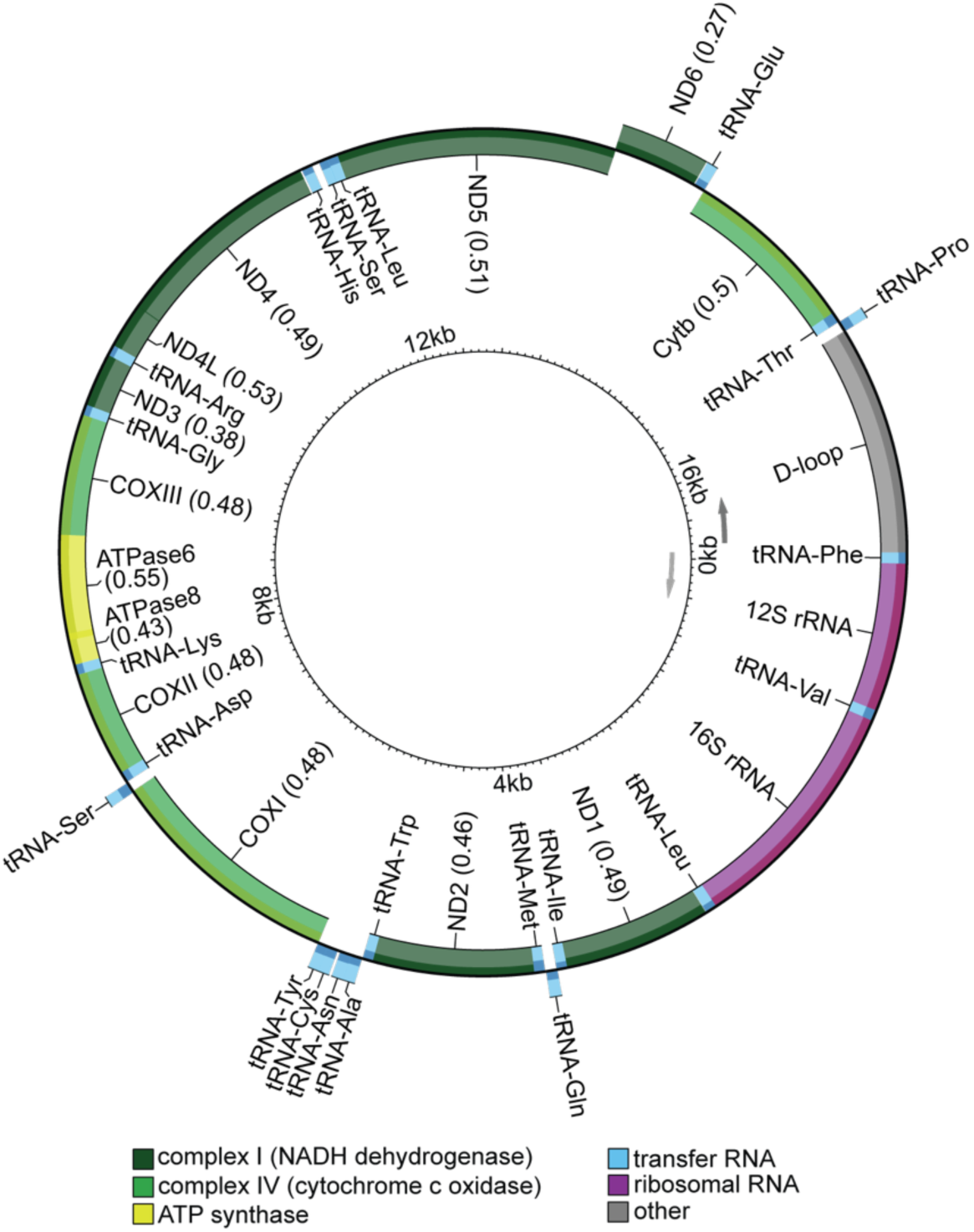
*Fukomys anselli* mitogenome annotation.

### Repetitive regions & transposable elements

Repeats and transposable elements were identified and annotated using Earl Grey (Baril et al. 2024). Repetitive elements span a total of 41.3% of the *F. anselli* genome, with retroelements (LINEs, SINEs, Penelope class, and LTR elements) constituting 35.6%, DNA transposons 3.3%, and simple repeats 1.0% (**Figure 3a**, **Table 3, Table S5, Figure S4**). The genomic fraction of repetitive regions is higher than that observed in *H. glaber* (28.9% in HetGla_female_1.0) or *F. damarensis* (30.3% in DMR_v1.0_HiC).

**Figure 3:**
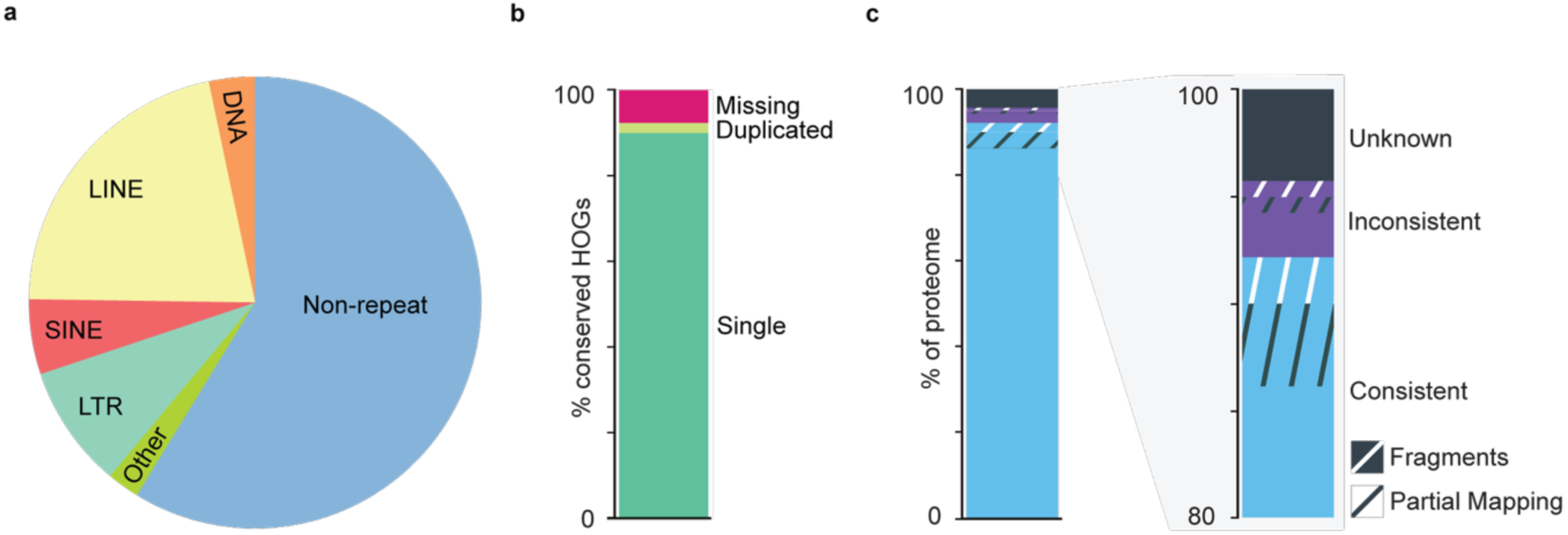
Repetitive regions and proteome. a) Proportion of the genome composed of repetitive regions and transposable elements. DNA = DNA transposon; LINE = long interspersed nuclear element; SINE = short interspersed nuclear element; LTR = long terminal repeat retrotransposon b) Proteome completeness. HOG = hierarchical orthologous gene c) Proteome consistency within the hystricomorph rodent hierarchical orthologous gene family.

**Table 3:** *Fukomys anselli* repetitive regions and transposable elements.

|  | Coverage (bp) | Copy Number | % Genome Coverage | TE Family Count |
| --- | --- | --- | --- | --- |
| **DNA** | 74,913,617 | 408,765 | 3.2942 | 397 |
| **Rolling Circle** | 140,571 | 673 | 0.0062 | 5 |
| **Penelope** | 9,165 | 48 | 0.0004 | 1 |
| **LINE** | 488,312,729 | 1,503,433 | 21.4727 | 265 |
| **SINE** | 121,651,153 | 1,137,637 | 5.3494 | 58 |
| **LTR** | 200,486,946 | 748,964 | 8.8161 | 593 |
| **Other (Simple Repeat, Microsatellite, RNA)** | 27,836,643 | 601,114 | 1.2241 | 12,474 |
| **Unclassified** | 24,898,076 | 173,619 | 1.0949 | 156 |
| **Total Repeats** | 938,248,900 | - | 41.2579 | - |
| **Non-Repeat** | 1,335,857,717 | - | 58.7421 | - |

Our assembly of *F. anselli* used both short read and ONT long-read sequencing; in contrast, both the *H. glaber* and *F. damarensis* genomes (HetGla_female_1.0; DMR_v1.0_HiC) were assembled using only short-read whole-genome sequencing, which can fail to capture the full extent of the repetitive genome (Lewis et al. 2016; Platt et al. 2016). We therefore compared our genome to a recent naked mole-rat genome (mHetGlaV3) assembled using similar long-read sequencing data. Indeed, this naked mole-rat genome reports a repetitive element composition strikingly similar to our findings in *F. anselli*: cumulatively 40.4% of the genome, with 36.2%, 2.6%, and 1.0% retroelements, DNA transposons, and simple repeats, respectively (Sokolowski et al. 2024).

### Gene structure annotation

Protein-coding regions of the curated and repeat-softmasked *F. anselli* assembly were annotated using BRAKER (Hoff et al. 2019; Gabriel et al. 2024). We generated RNA-seq from nine different tissues (liver, spleen, testes, calf muscle, kidney, heart, lung, brain, back skin) and combined them with protein sequences annotated in closely and distantly related species (**Table S2**). In total, we identified 29,094 transcripts across 18,256 predicted genes.

Assessment of proteome completeness via OMArk (Nevers et al. 2025) affirmed that *F. anselli* belongs to the hystricomorph rodent hierarchical orthologous gene (HOG) family. The *F. anselli* annotated proteome contains 92.35% of the 13,570 conserved Hystricomorpha HOGs and no contaminant sequences (**Table 4**, **Figure 3bc**). Further, among the 18,256 *F. anselli* predicted protein-coding genes, 92.18% are consistent with known gene families within Hystricomorpha; only 4.23% do not correspond to proteins in a known gene family (**Table 4**, **Figure 3c**). The species calculated to be most consistent with the HOGs in the annotated genome is *F. damarensis*, with 95.77% concordance; this estimate provides validation of phylogenetic proximity of these two species.

**Table 4:** *Fukomys anselli* proteome completeness and consistency.

| Completeness |  |  |
| --- | --- | --- |
| Conserved HOGs | 13570 (clade used: Hystricomorpha) |  |
| Single | 12210 (89.98%) |  |
| Duplicated | 322 (2.37%) |  |
|  | Duplicated, Unexpected | 279 (2.06%) |
|  | Duplicated, Expected | 43 (0.32%) |
| Missing | 1038 (7.65%) |  |
| Consistency |  |  |
| Total proteins | 18256 |  |
| Consistent | 16828 (92.18%) |  |
|  | Consistent, partial hits | 697 (3.82%) |
|  | Consistent, fragmented | 390 (2.14%) |
| Inconsistent | 656 (3.59%) |  |
|  | Inconsistent, partial hits | 136 (0.74%) |
|  | Inconsistent, fragmented | 131 (0.72%) |
| Contaminants | 0 (0.00%) |  |
| Unknown | 772 (4.23%) |  |

### Ongoing discussion of phylogenetic placement

There has been a long-standing debate regarding the taxonomic placement of several *Fukomys* species, including *F. anselli* (Faulkes et al. 2004; Ingram et al. 2004; Van Daele et al. 2007; Faulkes et al. 2017; Šumbera et al. 2023). Phenotypic and genotypic considerations may lead to re-designation of *F. anselli* (personal communication from Radim Šumbera). Regardless of potentially altered nomenclature, this genome assembly is utilizable for investigating *Fukomys* species with the karyotype of 2n=68 as well as closely-related species and hybrid animals.

In summary, our *de novo* genome of the Ansell’s mole-rat demonstrates chromosome-level contiguity and near-perfect gene completeness. We provide a novel annotated mitogenome, well-resolved repetitive regions and transposable elements, and a thorough protein-coding transcript annotation derived from nine diverse tissues. Given its completeness and quality, the *F. anselli* genome is comparable or superior to other available rodent genomes, providing a powerful resource to study the organismal phenotypes of Ansell’s mole-rat and resolving Bathyergidae phylogeny.

## DATA AVAILABILITY STATEMENT

The genome assembly and raw sequencing reads (WGS of ONT long reads and Illumina short reads, Hi-C, and RNA-seq) generated in this study are deposited at NCBI under the BioProject PRJNA1240209. The nucleotide sequences of the Ansell’s mole-rat nuclear and mitochondrial genomes assembled in this study are available at GenBank under the accessions (accession number pending) and PV670003, respectively. The gene structure and repeat annotation files associated with the Ansell’s mole-rat genome assembly are available on Zenodo (https://doi.org/10.5281/zenodo.15350947). The code used to process, assemble, and annotate the Ansell’s mole-rat genome is also available on Zenodo (https://doi.org/10.5281/zenodo.15489638).

Nuclear and mitochondrial genome sequences of other mole-rat species used in this study are publicly available and include: Damaraland mole-rat genome assembly DMR_v1.0_HiC (NCBI RefSeq: GCF_012274545.1) (Fang et al. 2014) and mitochondrion (NCBI RefSeq: NC_027742.1); naked mole-rat genome assemblies HetGla_female_1.0 (NCBI RefSeq: GCF_000247695.1) (Keane et al. 2014) and mHetGlaV3 (ENA: GCA_964261345) (Sokolowski et al. 2024), and mitochondrion (NCBI RefSeq: NC_015112.1). Proteomes used in this study are publicly available, and their corresponding accession numbers are listed in Supplementary Table S2.

## AUTHOR CONTRIBUTIONS (alphabetical)

Conceptualization– MB, VFB

Computational analysis– RC, VFB

Data generation– MB, MBehm, REW

Data curation– RC

Funding acquisition– AG, DTO, MF

Tissue collection– MBehm, SB

Animal maintenance– SB

Supervision– AG, DTO, MF

Visualization– RC, VFB

Writing (original draft)– MB, VFB

Writing (review and editing)– DTO, MB, RC, VFB

## Supporting information

Supplementary Figures and List of Supplementary Tables

## ACKNOWLEDGMENTS

We are thankful to Celine Reifenberg, Daniela Sohn, Marie-Luise Koch, Stefania Del Prete and Liana Penteskoufi for their support with the animal experiments. We thank the NGS Core Facility, German Cancer Research Center (DKFZ), for providing sequencing services.

