## Supplementary Figures and List of Supplementary Tables for "De novo genome assembly of Ansell’s mole-rat (*Fukomys anselli*)"

*Co-first author

**Content: 4 Supplementary Figures; List of Supplementary Tables (separate file)**

### Supplementary Figures

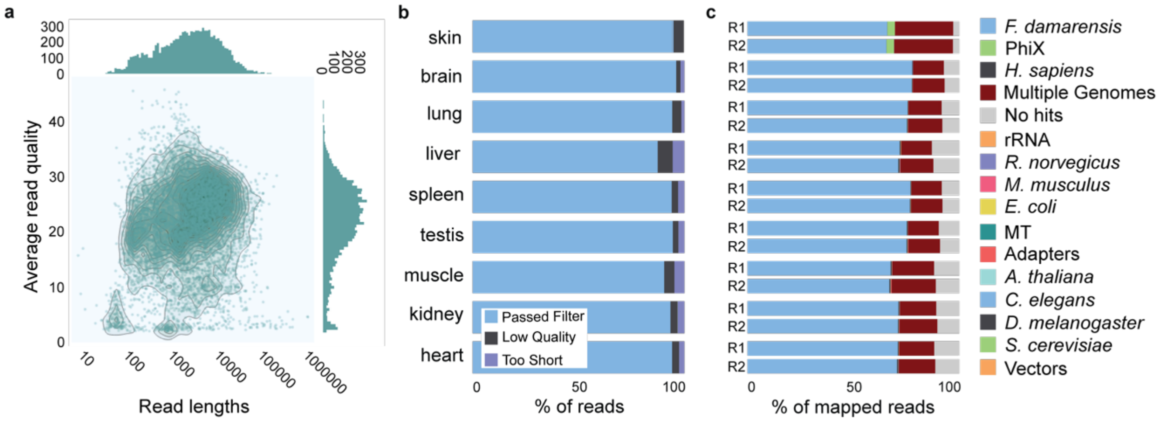


**Figure S1**: Sequencing quality control metrics. a) Pooled nanopore sequencing length and quality, per read. RNA-seq read quality (b) and predicted genome alignment (c) for nine tissues.

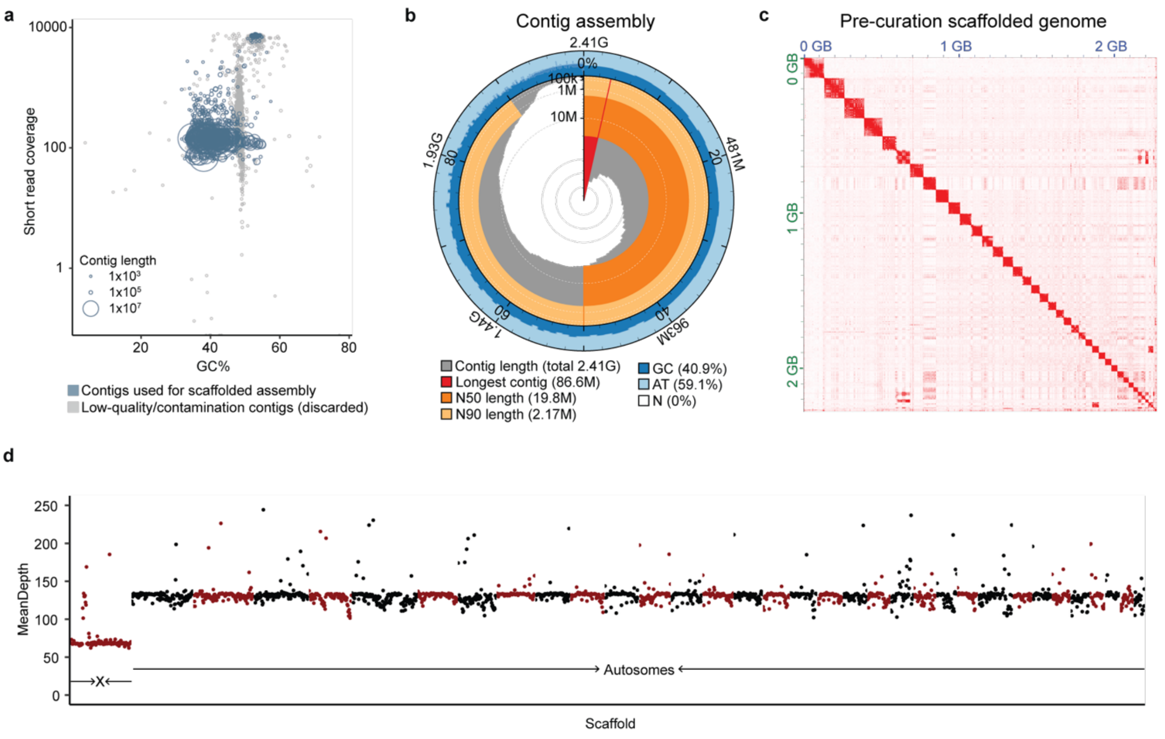


**Figure S2**: Intermediate steps in assembling the *Fukomys anselli* genome. a) Contig length, coverage, and GC content among assembled contigs. Grey contigs were identified as low-quality or contamination and so were not included in the final scaffolded assembly. b) Snail plot depicting the length and GC content of all contigs following purging of contamination contigs. c) Hi-C contact map of scaffolded genome prior to curation. The saturation of red corresponds to the number of contacts. d) Hi-C read coverage of the curated assembly along 1Mb bins. The average read depth is truncated at 250x coverage.

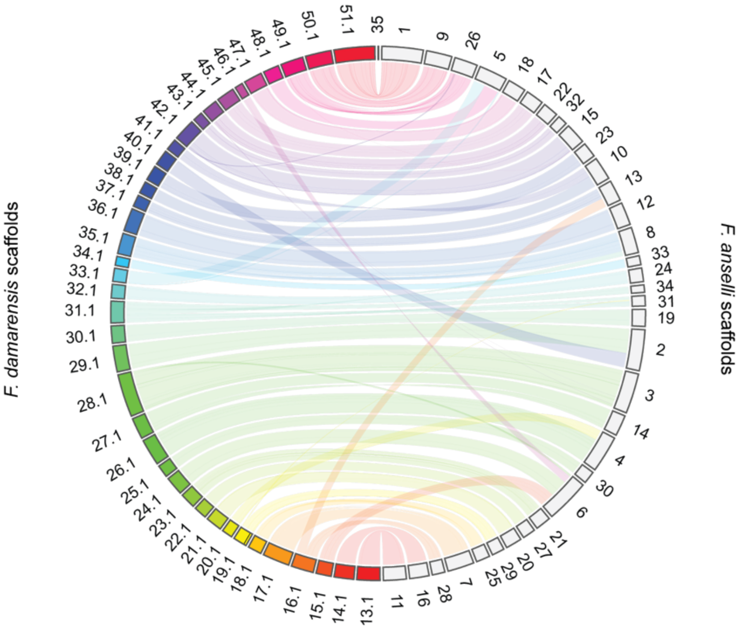


**Figure S3**: Circos plot depicting synteny between *Fukomys damarensis* (rainbow colored boxes) and *Fukomys anselli* (grey boxes) scaffolds for all *F. damarensis* scaffolds longer than 10Mb and for *F. anselli* scaffolds constituting 99% of the genome. The number of each scaffold is indicated, and *F. damarensis* scaffold names have been truncated to exclude the prefix "NW0229009".

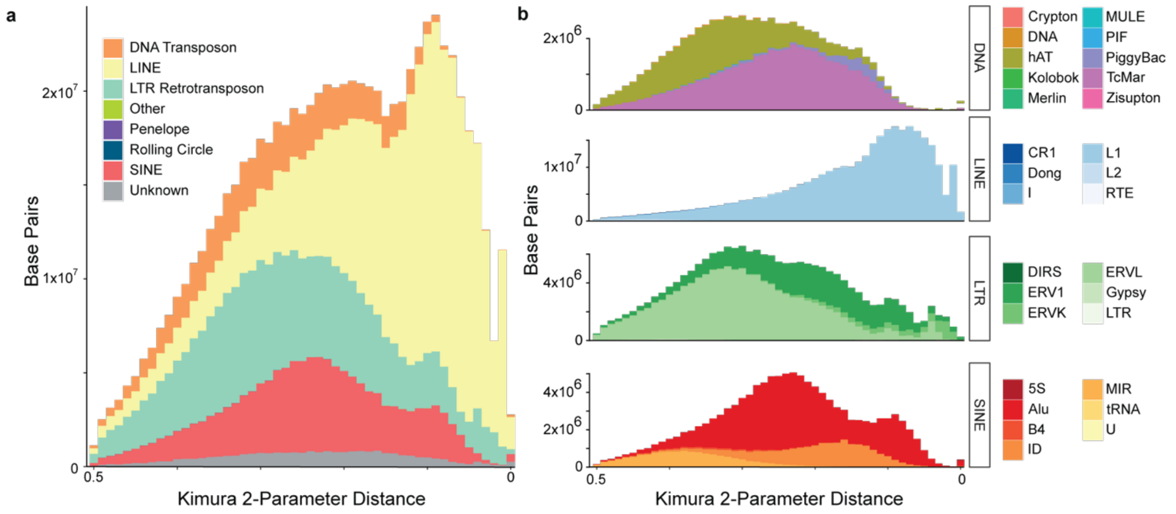


**Figure S4**: Kimura 2-parameter distance of transposable elements for all pooled (a) and family-resolved (b) classes in the *Fukomys anselli* genome. Some transposable element populations represented in the legends have too few occurrences to visualize at this resolution. DNA = DNA transposon; LINE = long interspersed nuclear element; SINE = short interspersed nuclear element; LTR = long terminal repeat

**Supplementary tables**

Table S1: Nanopore QC statistics per run

Table S2: List of NCBI annotated rodent proteomes used for annotation

Table S3: Assembly statistics for intermediate *Fukomys anselli* genome assemblies

Table S4: Number of informative Hi-C reads across contigs

Table S5: Family-level repetitive regions and transposable elements
